# Single-cell quantitative analysis of skeletal muscle cell population dynamics during regeneration and ageing

**DOI:** 10.1101/222158

**Authors:** Lucia Lisa Petrilli, Filomena Spada, Claudia Fuoco, Elisa Micarelli, Alessio Reggio, Marco Rosina, Cesare Gargioli, Luisa Castagnoli, Gianni Cesareni

**Affiliations:** Department of Biology, University of Rome Tor Vergata, Rome, Italy

## Abstract

The skeletal muscle is populated by a variety of different mononuclear cell types that actively contribute to tissue homeostasis. When the tissue is stressed by exercise or by an acute or chronic insult, the different cell types are activated, exchange signals and initiate a finely-orchestrated regeneration process to prevent the loss of muscle mass. This cell variety is exacerbated by an additional intra population heterogeneity, where the cell population boundaries often lose their significance. Here, we applied a high-dimensional single-cell mass cytometry analysis to solve the cellular and molecular complexity of the muscle tissue in different physiological and pathological conditions. Taken together, our results provide a comprehensive picture both of muscle cell population homeostasis during ageing and of the changes induced by a perturbation of the system, be it chronic (dystrophy) or acute (cardiotoxin).

## Introduction

In physiological conditions, the adult skeletal muscle is characterized by a relatively low cell turnover^1^. However, in response to trauma, whether it be of mechanical or chemical origin or caused by genetic defects, the muscle tissue activates a finely orchestrated and efficient regeneration process, that partially recapitulates the embryonic development^2^.

The dissolution of the myofiber plasma membrane, triggered by the damage, induces an influx of calcium ions causing the activation of calcium-dependent proteases^3^. This in turn results in myofiber necrosis that causes an inflammatory response initiated by the circulating neutrophils moving into the injury site. Neutrophils release pro-inflammatory cytokines that attract macrophages, entering the damaged muscle pro-inflammatory environment^4,5^. The participation of macrophages in this process is commonly divided into an early wave of pro-inflammatory macrophages (M1), involved in the removal of the tissue necrotic debris, which is followed by a phenotype switch, called macrophage polarization, leading to a second wave of antiinflammatory macrophages (M2). M2 macrophages secrete cytokines that trigger the activation and differentiation of satellite cells (SCs), a stem cell population with myogenic potential^6^.

Satellite cells, located beneath the myofiber basal lamina, count for the 2–5% of the sublaminar nuclei and are the central players in skeletal muscle tissue regeneration^7,8^. Once activated, following exercise, injury or in degenerative diseases, satellite cells escape their mitotically quiescent state and re-enter the cell cycle. Finally they proliferate and activate a terminal differentiation program, characterized by changes in the expression of specific myogenic transcription factors, which culminate in the fusion with damaged myofibers or in the formation of new myotubes^9^. A small fraction of the activated satellite cells go through an asymmetric division where one of the daughter cells goes on to amplify the pool of myogenic cells while the other returns to quiescence thus contributing to maintain muscle tissue regenerative potential by replenishing the reserve of satellite stem cells^10,11^. This complex process ensures an efficient tissue repair in a healthy muscle. With ageing, satellite cells progressively lose their self-renewal and regenerative capacities and the regeneration process becomes less efficient^12–14^.

In addition to the central role of the interplay between monocyte/macrophages and SCs in muscle tissue healing, the efficient structural and functional regeneration of muscle tissue integrity requires a well-coordinated crosstalk between additional muscle resident mononuclear cell types. These functionally diverse cell populations are characterized by the expression of specific surface antigens and modulate the regeneration process by sending pro or anti differentiation signals to progenitor cells.

Among these cell types, which directly or indirectly contribute to maintain myofiber homeostasis, a population of interstitial cells, called fibroadipogenic progenitors (FAPs), has recently gained attention for its prominent role in muscle regeneration^15^. In resting conditions, muscle myofibers negatively control FAP adipogenic differentiation^16^. After acute injury, FAPs are activated by the cytokines IL-4 and IL-13, secreted by eosinophils during the early regeneration phases^17^ and rapidly expand to produce paracrine factors (i.e. IL-6) that stimulate satellite-cell mediated regeneration. In addition FAPs, during the regeneration process, contribute to the reassembly of the muscle tissue architecture by depositing extracellular matrix (ECM) proteins^16–18^. Towards the completion of the regeneration process, the excessive FAPs, generated during the expansion phase, are removed by macrophage-promoted apoptosis while the remaining ones return to the initial quiescent state^19^. In contrast, in chronic injury caused by genetic disorders, as in muscle dystrophies, the change in the inflammatory milieu counteracts FAP apoptosis causing their accumulation and differentiation as fat and fibrotic deposits that disrupt the muscle architecture^16,19^.

Understanding the details of this finely orchestrated process has been the object of several studies over the past years^15,16^. However, we are still missing an integrated quantitative picture of the cell population dynamics after acute or chronic damage. In addition, most of the studies are based on the analysis of bulk cell populations defined by the expression of a couple of specific markers. Little is known about the heterogeneity of these populations and how they change in time when the muscle physiology is perturbed. These questions can only be addressed by approaches that are based on single-cell analysis. For this reason, here we apply high-dimensional single-cell mass cytometry to describe the dynamic changes in the muscle cell populations during ageing and after acute or chronic damage^20,21^.

## Results

### Changes in the cell population abundance in ageing wild type and *mdx* mice

We first investigated the changes caused by ageing in mononuclear cell population abundance in mouse muscles at four different time points corresponding to young (1.5 months), mature (3.5 months), middle age (8 months) and old (16 months). In addition to wild type C57BL/6, we also considered a mouse model (*mdx*) of Duchenne Muscular Dystrophy. The genome of this mouse carries a loss-of-function mutation in the dystrophin gene, condition that causes muscle fragility and chronic inflammation^22^. Histological sections of the tibialis anterior (TA) muscle of ageing wild type mice after hematoxylin and eosin (H&E) or Sirius red staining did not reveal any macroscopic change (Fig. 1a, upper panel). Myofibers of relatively uniform size and cross sectional area, with nuclei located at the periphery of the muscle cell and a thin layer of connective tissue between fibers, characterize muscles of wild type mice. In addition, Sirius red staining did not identify a significant increase in collagen deposition with ageing (Fig. 1b, upper panel).

**Figure 1.**
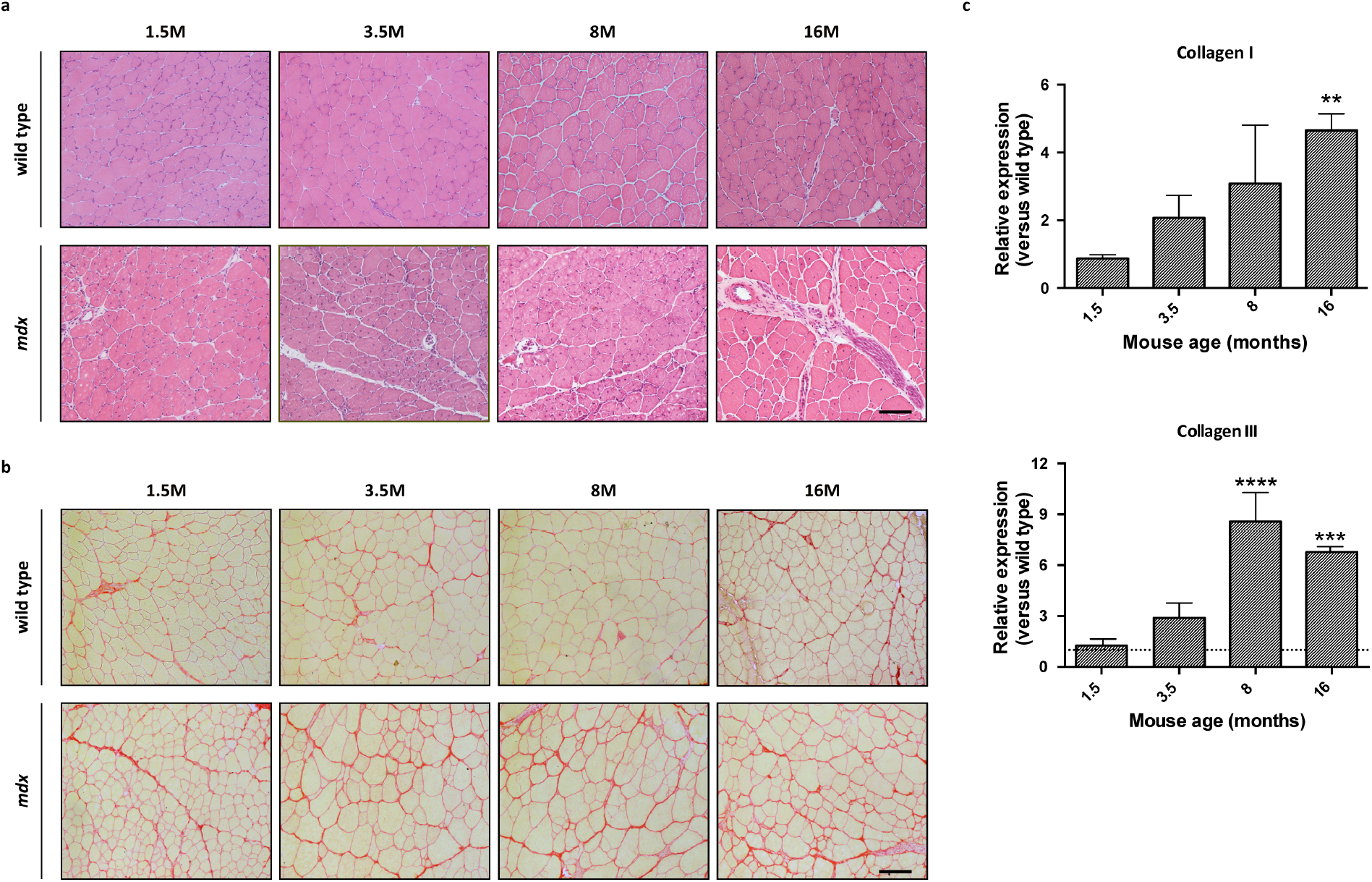
Effect of ageing on wild type and *mdx* skeletal muscle. Representative images of **(a)** hematoxylin and eosin and **(b)** Sirius red histological staining on the *tibialis anterior* muscle of wild type and *mdx* mice of different ages (1.5, 3.5, 8 and 16 months). **(c)** RT-qPCR analysis of the expression level of *Collagen I* and *Collagen III* mRNA in *mdx* TA muscles at the indicated ages. Values are mean ± SEM and are expressed as fold change compared to the wild type expression. β-Actin was used as control gene (n = 3; ** p < 0.01; *** p < 0.001; **** p < 0.0001 wild type vs *mdx*). Scale bars (**a, b**): 100 μm.

By contrast, *mdx* TA are characterized by the presence of centrally nucleated regenerating myofibers of different shape (Fig. 1a, lower panel). Moreover, significant fibrotic deposition was highlighted in older *mdx* mice and confirmed by Sirius Red staining (Figure 1b, lower panel). Further, comparative qPCR experiments proved the proportional and time-dependent increase in the *Collagen* I (Coll-1a2) and *Collagen* III (Coll-3a1) mRNA in *mdx* mice (Figure 1c).

To monitor the changes in the relative abundance of the different mononuclear cell populations caused by the ageing process, we assembled a collection of 23 metal-tagged antibodies (Supplementary Table 1). The antibody panel chosen for the study was composed by 15 metal-tagged antibodies that were specific for antigens expressed by several populations that constitute the muscle niche, as well as by the hematopoietic compartment. Furthermore, the analysis was enriched by including 8 additional antibodies, raised against intracellular antigens, used to monitor the activation of specific pathways in the cell types identified by their antigenic signature.

Single-cell suspensions of hind limb muscles were isolated from different age wild type and *mdx* mice. The number of mononuclear cells was basically higher, by a factor of 1.5, in muscles from *mdx* mice (Supplementary Fig. 3a). The isolated mononuclear cells were further processed for mass cytometry (Fig. 2a). To obtain comparable readouts a barcoding protocol^23^ was used before labeling the different samples with the panel of antibodies in a single test tube. Reagents against the FoxP3, Pan-Actin, interleukin-6, phospho-Akt, Cleaved Caspase3, CD4 and phospho-Erk antigens gave a non-modal intensity distribution (Supplementary Fig. 1). Seven readouts (CD45, CD34, Sca-1, α7-integrin, F4/80, CD90.2, CD206) displayed a bimodal distribution with two clear peaks of cells expressing different levels of the signal while the remaining were characterized by a broader reactivity distribution often containing several peaks (Supplementary Fig. 1). Figure 2c reports some examples of two-dimensional scatter plots, revealing cell sub-populations that are either characteristics of one of the two mouse model systems and differences over the considered time frame. Additional scatter plots are reported in Supplementary Fig. 2.

**Figure 2.**
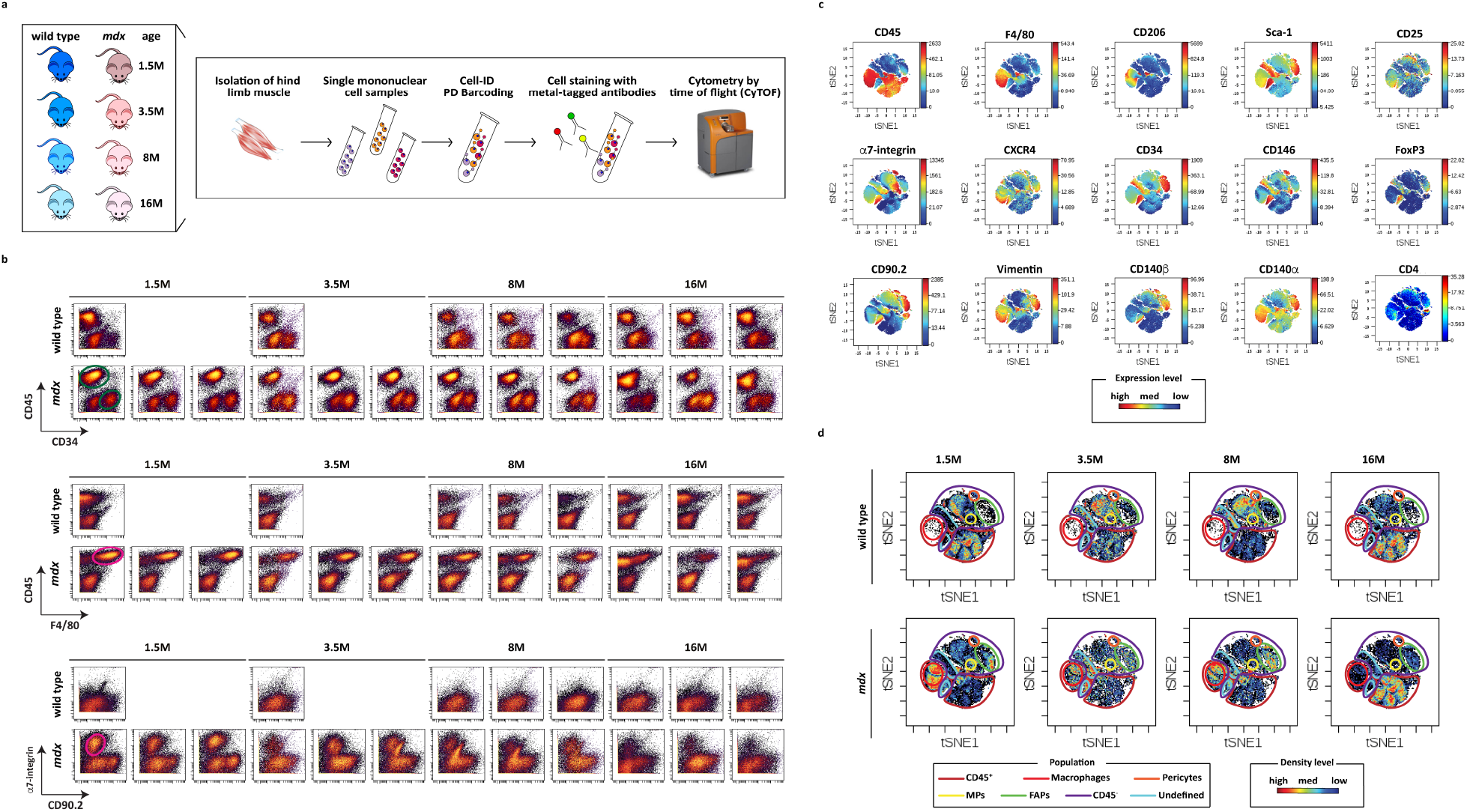
CyTOF2 analysis on different age wild type and *mdx* muscles. **(a)** Mass cytometry workflow. Hind limb muscles were collected from wild type and *mdx* mice of different ages and mononuclear cells were dissociated from the myofiber by enzymatic digestion (DNase I, collagenase A and dispase II). The obtained cell suspension was next subjected to sequential filtrations. The resulting single mononuclear cell samples from the different conditions were “barcoded” with a specific combination of three different palladium isotopes and then combined in a single sample that was subsequently labeled with heavy metal coupled antibodies specific for relevant cell antigens. The labeled cells were finally analyzed by the CyTOF2 mass cytometer instrument. **(b)** Scatter plots of some of the antibodies that displayed a bimodal intensity distribution in the ageing experiments. **(c)** viSNE maps of mononuclear cell populations of 1.5 month old *mdx* mice. The different maps are colored according to the expression levels of the selected antigens (red, high expression; blue, low expression). **(d)** Representative viSNE dot-plot maps showing the density distributions (red, high density; blue, low density) of mononuclear cells from muscles of wild type or *mdx* mice of different ages. According to the expression profile, we highlighted different populations that are relatively homogeneous in their expression profiles and, where possible, we labeled them with the name of the cell type better matching their profile according to literature.

To make full use of the multi-parametric data and to identify the cell populations of interest, the mass cytometry dataset was analyzed by applying the viSNE algorithm. Figure 2c shows a representative viSNE analysis output. Each map represents a bi-dimensional projection of the n-dimensional investigated space. Each dot is characterized by a specific position in the plot, defined by tSNE coordinates (tSNE1/tSNE2), and corresponds to a single-cell. The used color can vary from red (high expression) to blue (low expression) according to the expression level of the chosen antigen.

By looking at the expression of the different markers in the viSNE plot, and by comparing it with literature information, we were able to assign each area of the plot to a specific cell type (circled in Fig. 2d) (Supplementary Table 2).

Note that because of the absence of suitable reagents, we were not able to selectively identify all the cell populations of interest and some events still remained undefined. For instance, we could not identify a good reagent specific for type 1 macrophages.

The visual inspection of the colors (densities) of the different areas of the plots revealed some changes in the population relative abundance. For example, while the macrophage population (F4/80^+^) remained low and did not vary significantly with age in wild type mice, the muscles of *mdx* mice were colonized by a more numerous macrophage population that became less abundant with time (Fig. 2d).

#### CD45^+^ hematopoietic cells

An initial analysis of mononuclear cells, carried out by gating for the conventional hematopoietic marker CD45, highlighted that the 50/50 ratio between CD45^−^/CD45^+^ cells was maintained without significant variations during the considered life-span, with an exception (Fig. 3a). In fact, an important difference in the CD45 ratio was observed in 8 month old wild type mice.

**Figure 3.**
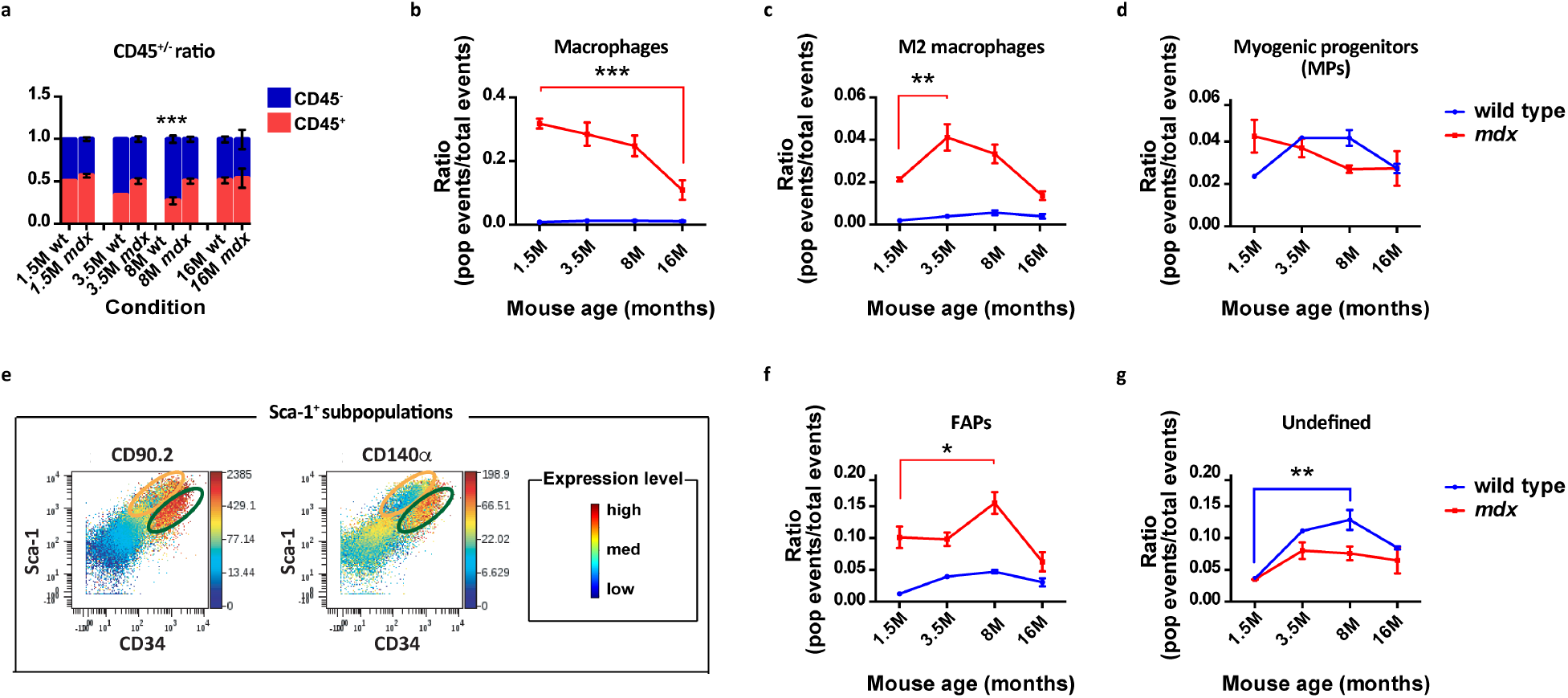
Dynamics of muscle monuclear cell population in ageing wild type and *mdx* skeletal muscle. **(a)** The bar plots represent the relative abundance of CD45^−^ (blue) and CD45^+^ (red) muscle mononuclear cells in the different conditions tested (*** *p* < 0.001 CD45^+^ *vs* CD45^−^ in 8 month old wild type mice). **(b-d)** The graphs illustrate the dynamics of muscle mononuclear populations, isolated from the hind limb muscles of different age wild type (blue) and *mdx* (red) mice. The trend of **(b,c)** macrophages and of the **(d)** myogenic progenitors during ageing was clearly outlined (**b**, *** *p* < 0.001 1.5 *vs* 16 month old *mdx* mice; **c**, ** *p* < 0.01 1.5 *vs* 3.5 month old *mdx* mice). **(e)** Representative scatter plots highlighting the existence of two Sca-1/CD34 positive subpopulations showing a different expression for CD90.2 (right) and CD140α (left) (red, high expression; blue, low expression). **(f,g)** The graphs illustrate the abundance of FAPs and of an undefined population during the ageing process (**f**, * *p* < 0.05 1.5 *vs* 8 month old *mdx* mice; **g**, * *p* < 0.05 1.5 *vs* 8 month old wild type mice). All the diagrams were generated by collecting data (n=1-3) by mass cytometry and by analyzing them through the viSNE algorithm. The ratio represents the fraction of cells expressing the given antigens in the total mononuclear cell population. Statistical significance was evaluated by a twoway ANOVA.

A larger significant difference between the two models was observed when considering the abundance of F4/80^+^ cells, including eosinophils, M1 and M2 macrophages and monocytes. These cell types play an important role in the chronic inflammatory phenotype of dystrophic muscles.

In particular, *mdx* F4/80^+^ cell number was relatively high and constant until 8 months after birth and significantly higher, almost thirty-fold, than the wild type levels (Fig. 3b). In addition, it decreased, but still remained higher than wild type, in the older *mdx* mice (16 months) when it reached the abundance of 10% of the total events, reflecting the attenuation of the inflammatory response in the old *mdx* mice. By contrast, the wild type macrophage cell number was very low (~1%) along the monitored time span, consistent with the absence of inflammatory stimuli in physiological conditions.

We also investigated the abundance of F4/80 populations expressing CD206, expressed in M2 macrophages. This analysis revealed that also the M2 macrophage number was higher if compared to wild type. Figure 3c shows the trend of the M2 populations in muscles of wild type and *mdx* mice of different ages and reveals that, in *mdx* mice, the anti-inflammatory M2 macrophages are lower in the regeneration windows of 1.5 and 8 months (2 and 3% respectively), while they reach a peak during the degeneration phase of 3.5 months (4%). The observed decrease of M2 macrophages in old *mdx* mice confirms the decline of the inflammatory process as the pathology progresses.

#### Sca-1 cell populations

To well characterize the nature of muscle residing cells as well as to define their dynamics in time, we next considered the CD45^−^ compartment, including the non-hematopoietic muscle cell types. Therefore, we first analyzed the cell populations expressing low levels of the Stem cell antigen (Sca-1).

Among Sca-1^−^/CD45^−^ cells, we first considered the myogenic progenitors (MPs), satellite cells and myoblasts, identified as α7-integrin^+^, CXCR4^+^ and CD34^L^ cells. We did not observe any significant difference between *mdx* and wild type at all considered time points. (Fig. 3d). Thus, we concluded that α7-integrin^+^ cell abundance, comprising satellite cells and myoblasts, remained constant in time (3–4 % range) in both wild type and *mdx* mice.

Focusing on Sca-1^+^ cells, we observed two large populations. They both express the CD34 marker, but show different levels of CD140α and CD90.2 (Fig.3e). The CD140α^+^ and CD90.2^+^ population (circled in green) has an expression profile that is compatible with that of the fibroadipogenic progenitors (FAPs). This population includes approximately 10% of the total muscle mononuclear cells in young *mdx* mice (1.5 and 3.5 months) and, after reaching its peak in 8 month old *mdx* mice (15% of total events), it decreased in the old mice (Fig. 3f). On the contrary, the medium fraction of FAPs extracted from wild type mice was stable around 3.5%.

The remaining Sca-1^+^/CD34^+^ and CD140α^−^/CD90.2^−^ population (circled in yellow in Fig. 3e) showed significative variation only in the *mdx* mice aged 8 months and no relevant increase at the other time points in both wild type and *mdx* mice (Fig. 3g).

#### CXCR4 positive α7-integrin^+^ cells

Despite the fact that a strong variation was not observed in the number of MPs (Fig.3d), the high dimensional analysis made possible by mass cytometry, revealed that wild type and mdx α7-integrin^+^ cells differ in the expression of the CXCR4 antigen. CXCR4 is the receptor for the chemokine SDF-1 (Stromal Derived Factor-1) which is secreted in the muscle tissue in response to many cell pathways such as chemotaxis, proliferation or differentiation^24^. In particular, we observed that the α7-integrin^+^ cells derived from *mdx* muscles contained a higher fraction of cells expressing the CXCR4 antigen, especially in the youngest *mdx* mice (Fig. 4a). This observation lead us to speculate that the SDF-1/CXCR4 crosstalk may be involved in the dystrophic pathology.

**Figure 4.**
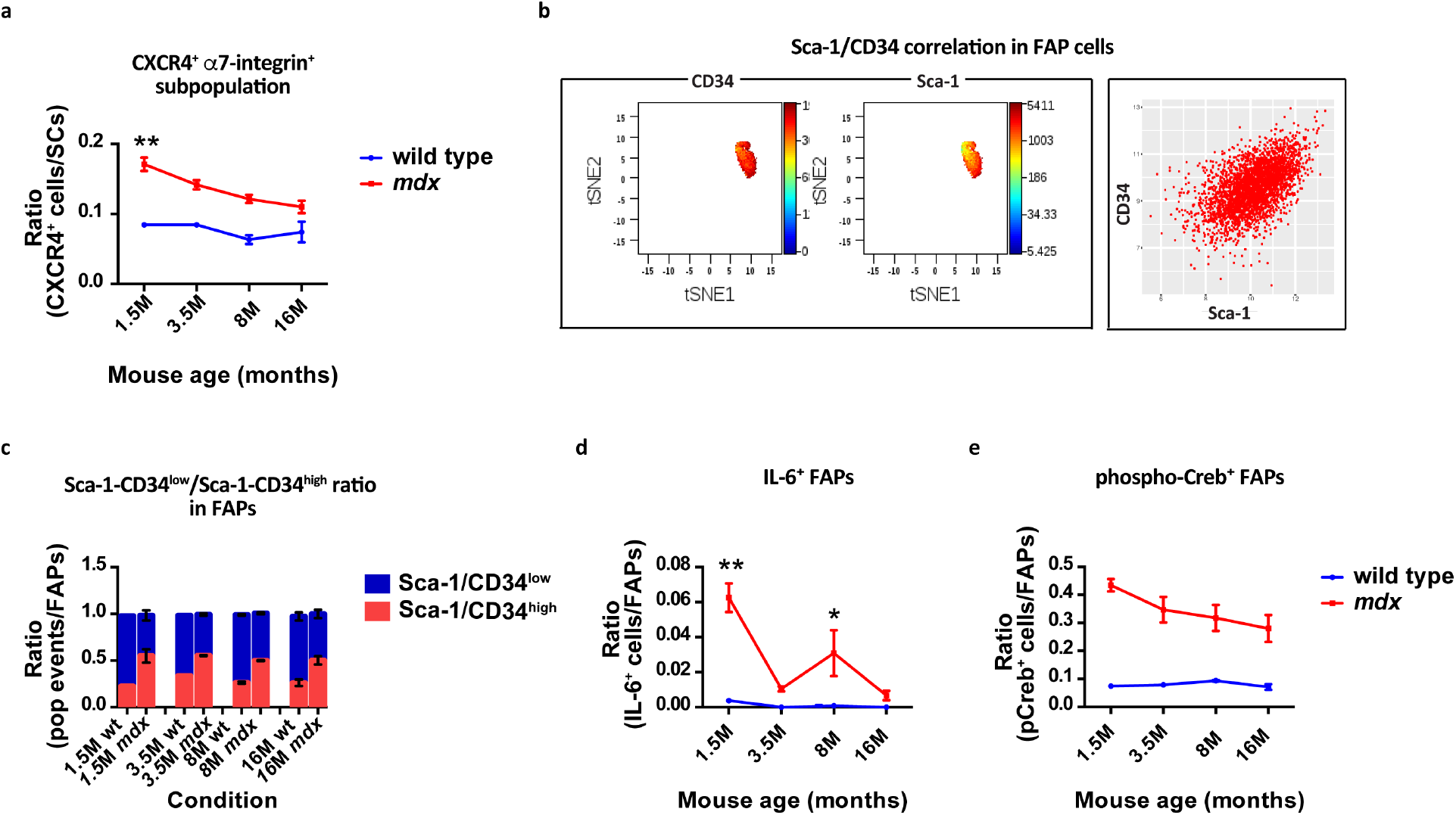
Satellite and fibroadipogenic populations during wild type and *mdx* tissue ageing. **(a)** Fraction of α7-integrin^+^ cells expressing the CXCR4 receptor (** *p* < 0.01 wild type vs *mdx* 1.5 month old mice). **(b)** viSNE density distribution (left) and scatter plot (right) graphs showing SCA-1/CD34 marker correlation in FAPs from 8 month old *mdx* mice. **(c)** Bar plots representing the relative abundance of Sca-1/CD34^low^ (blue) and Sca-1/CD34^high^ (red) FAPs in the different conditions tested. **(d-e)** Fraction of FAP cells expressing the phospho-Creb and IL-6 antigens during ageing of wild type and *mdx* mice (**d**, * *p* < 0.05 wild type *vs mdx* 8 month old mice; ** *p* < 0.01 wild type *vs mdx* 1.5 month old mice). The diagrams were generated by collecting data (n=1-3) by mass cytometry and by analyzing them through the viSNE algorithm.

#### Fibroadipogenic populations

When in-depth investigating the fibroadipogenic progenitor population, we observed that the expression of the self-renewal Sca-1 marker was associated to the expression of CD34 (Fig. 4b), a marker of cell proliferation. More specifically we saw that the cell events that were highly positive for Sca-1 also express high levels of CD34. For this reason, we evaluated the abundance of the Sca-1/CD34^high/low^ ratio in FAP cells from ageing wild type and *mdx* mice.

Interestingly, as shown in Fig. 4c, we observed that the number of cells expressing high levels of Sca-1 was higher in *mdx* mice in all the different analyzed conditions but, as in wild type mice, the ratio between Sca-1^L^CD34^L^/Sca-1^H^CD34^H^ expressing cells remained constant with ageing. This observation lead us to speculate that the relative abundance of the two populations could have an important role in ensuring tissue homeostasis.

Finally, the antibody panel that we used in the mass cytometry analysis also included antibodies against signaling proteins. Among them, remarkable results were obtained when considering Interleukin-6 (IL-6), one of the pro-myogenic factors secreted by FAPs to stimulate SC mediated regeneration^18^, and phospho-Creb, otherwise related to the activation of the adipogenic pathway^25^.

As shown, *mdx* FAP cells were positive for the expression of IL-6 and phosho-Creb (Fig.4 d, e), further confirming their double role in the degeneration of skeletal muscle tissue typical of the *mdx* mice.

Taken together these results confirmed a non-physiological persistence of FAPs within the dystrophic muscles that contributes to the pathological outcome.

### Muscle cell population dynamics following acute injury in wild type and *mdx* muscles

To gain further insight into skeletal muscle tissue, we next aimed at describing the dynamics of the previously characterized cell populations, following acute damage.

To this end we injected snake venom cardiotoxin (CTX) into the muscles of wild type and *mdx* mice (Fig. 5a). CTX is a protein kinase C specific inhibitor that, by inducing the depolarization and hypercontraction of muscle fibers, damages the structure of the cell membrane inducing cell death in a controlled and reproducible manner^26^.

**Figure 5.**
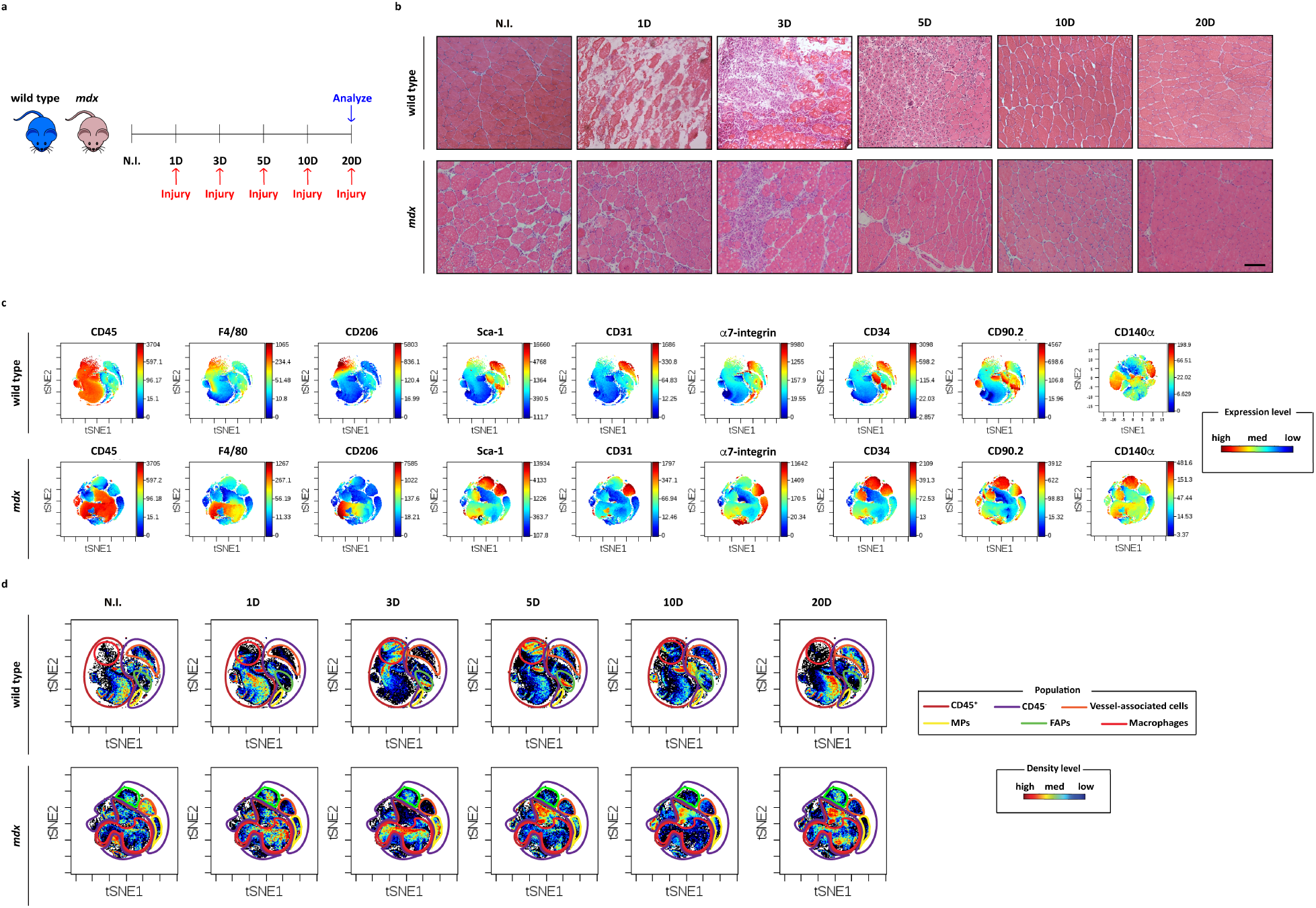
Cardiotoxin induced injury on wild type and *mdx* skeletal muscle tissue. **(a)** Scheme of the experimental procedure. Young, 1.5 months old, mice were injected intramuscularly with cardiotoxin (10 μM). At time 1,3,5,10 and 20 days after CTX injection, mice (3 for each time point) were sacrificed and the TA muscles collected for histological and mass cytometry analysis. **(b)** Representative images of hematoxylin and eosin histological staining on the tibialis anterior muscle of wild type and *mdx* mice transverse sections. Scale bar: 100 μm. **(c)** viSNE map examples of the multi-parametric mass cytometry analysis of mononuclear cells from 1 day post-injury wild type and *mdx* mice. Colors represent the expressions level of the selected antigens (red, high expression; blue, low expression). **(d)** Representative viSNE dot-plot maps showing the identified skeletal muscle niche populations. Colors map the cell density in the different areas of the plot (red, high expression; blue, low expression).

Figure 5b reports some representative histological sections of wild type and *mdx* tibialis anterior (TA) muscles injected with 10 μl of 10 μM cardiotoxin (CTX) and analyzed at different time points after the injury (1, 3, 5, 10 and 20 days). The non-injured (N.I.) wild type muscle sections are characterized by polygonal fibers, with limited interstitial space and containing peripheral nuclei.

The largest alterations of this typical muscle organization are observed at day 1 and 3 after cardiotoxin injury when the myofiber structure is disrupted and in the extended interstitial space necrotic areas and inflammatory infiltrates are clearly visible. A few days later, starting from day 5, the regeneration process enters its active phase and centrally nucleated myofibers appear. 20 days from the injury the physiological muscle architecture is fully restored.

In *mdx* injured muscles (Fig. 5b, lower panel), by contrast, already before the cardiotoxin insult we could observe inflammatory interstitial cells and centrally nucleated myofibers of different size. In addition, no significant histological differences were observed over the analyzed time frame.

Next, we were interested in describing the cell population dynamics during the regeneration process in the two systems. To this end, the hind limb muscles of wild type and *mdx* mice were collected and enzymatically digested to isolate mononuclear cells at different times after cardiotoxin treatment.

Supplementary Figure 3b represents the trend of the total number of cells, extracted from cardiotoxin injured wild type and *mdx* muscles. As shown in the graph representation, wild type mononuclear cells display a characteristic kinetic during the regeneration process. In particular, their number increases after the damage and reaches a peak at day 3. Starting at this time point, the number of mononuclear cells decreases, returning to almost baseline already at day 10. In the *mdx* mice by contrast, no significant fluctuations were observed after cardiotoxin injury since the total number of extracted mononuclear cells did not increase after injury and was very similar over all the regeneration period.

The mononuclear cell samples from both animal models were separately labeled by a barcoding protocol and analyzed by mass cytometry, with a panel of 23 metal-tagged antibodies (Supplementary Table 1) targeting antigens expressed in cells of the muscle and of the hematopoietic compartments.

By mapping the expression of selected surface antigens on the different areas of the plots (Fig. 5c), we were able to map on the viSNE plots the wild type and *mdx* cell populations participating in the skeletal muscle tissue response to cardiotoxin (Fig. 5d) (Supplementary Table 2). Figure 6 illustrates the dynamics of four selected populations.

**Figure 6.**
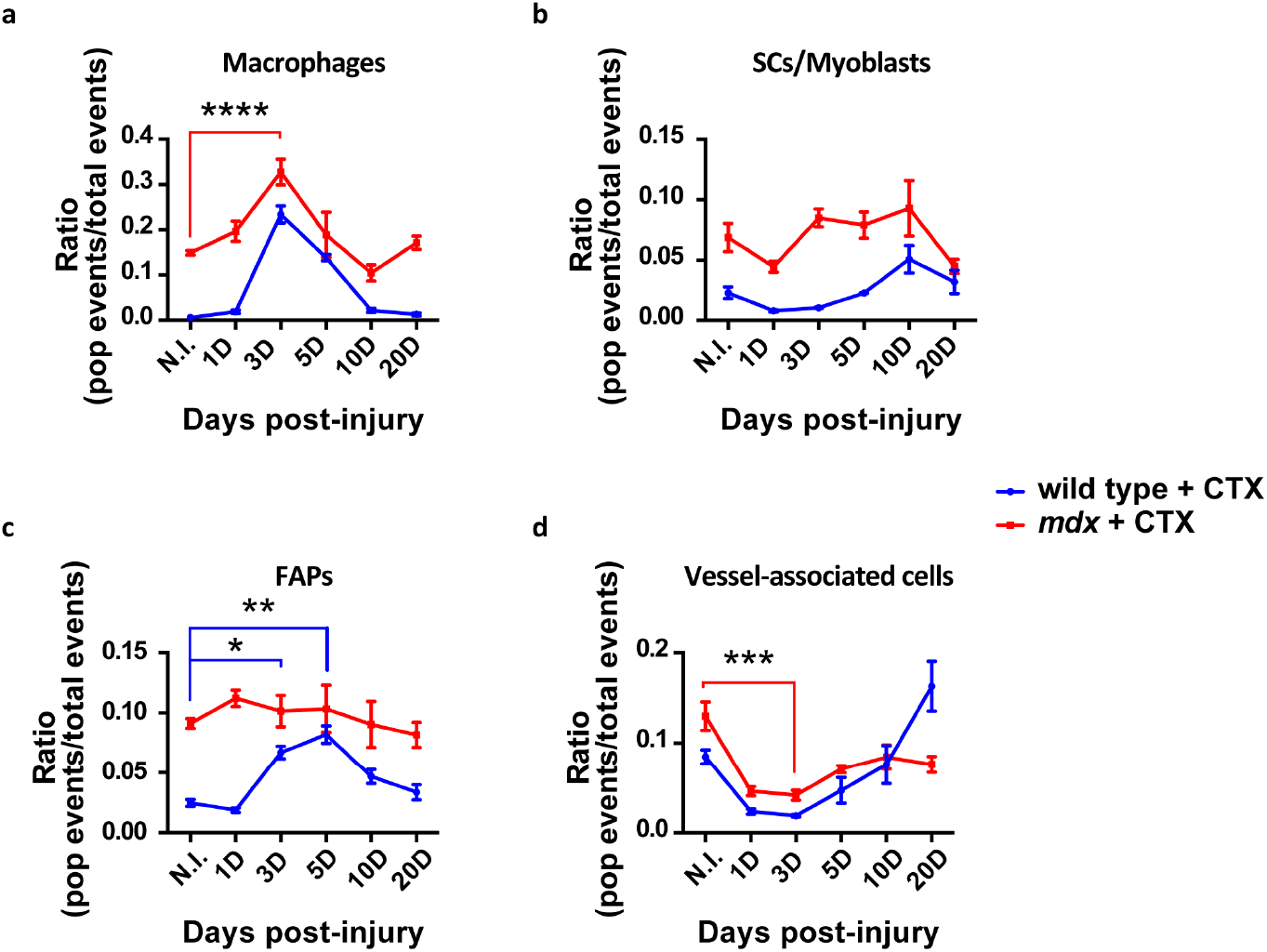
Dynamics of muscle monuclear cell populations after cardiotoxin-induced injury in wild type and *mdx* skeletal muscle. The graphs illustrate the dynamics of muscle mononuclear populations, isolated from the CTX-injected hind limb muscles of wild type (blue) and *mdx* (red) mice in two independent experiments: **(a)** Macrophages (F4/80^+^), **(b)** Satellite cells and myoblasts (α7-integrin^+^, CXCR4^+^) **(c)** fibroadipogenic progenitors (Sca-1^+^, CD140α^+^, CD34^+^, CD90.2^+^, Vimentin^+^) and **(d)** vessel-associated cells (CD31^+^, Sca-1^+^, CD34^+^, CD146^+^). The diagrams were generated by collecting data (n=3) by mass cytometry and by analyzing them by the viSNE algorithm. The ratio represents the fraction of cells expressing the given antigens over the total number of mononuclear cells. Statistical significance was evaluated by a two-way ANOVA (**a**, **** *p* < 0.0001 non injured, N.I. *vs* 3 day injured *mdx* mice; **c** * *p* < 0.05 non injured, N.I. *vs* 3 day injured wild type mice; ** *p* < 0.01 non injured, N.I. *vs* 5 day injured wild type mice; **d** *** *p* < 0.001 non injured, N.I. *vs* 3 day injured *mdx* mice).

We first focused on the CD45^+^/F4/80^+^ hematopoietic cells, since they have the important function of participating in the regeneration process with the double role of removing debris and triggering the repair machinery by stimulating the myogenic compartment to repair the damage.

In physiological conditions, the F4/80 positive hematopoietic cells, including eosinophils, monocytes and M1 and M2 macrophages, count for less than 1% of the total recorded events (Fig. 6a). Already 3 days after the injury (3D), however, their number increases, reaching a maximum of approximately 20%. From this time point on, their number starts to decrease and the difference with the non-injured (N.I.) control number gradually diminishes. Finally, the hematopoietic cell number returns to baseline levels at 20 days.

The hematopoietic population of the *mdx* mice is more numerous than wild type before injury (15%) (Fig. 6a), and similarly to wild type also shows significant changes following cardiotoxin treatment.

After being stimulated by a combination of signals sent from injured myofibers, blood vessel and inflammatory cells, the myogenic progenitors quickly activate to explicate their primary pro-regenerative role ^27^. Even the fibroadipogenic progenitors, upon stimulation, are known to exert their pro-myogenic function through the secretion of inflammatory cytokines^18^.

Given that, we first evaluated the myogenic progenitor (MP) dynamic of participation into the regeneration process.

An initial change of α7-integrin^+^ population, including satellite cells and myoblasts, was observed starting 24 hours after cardiotoxin injury in the wild type regeneration model. Once activated as a reaction to the cardiotoxin injury, their number slowly increases until reaching their maximum 10 days post-injury, accounting for about 5% of the total events recorded by mass cytometer. At 20 days, the α7-integrin^+^ cell number returned to baseline level. The *mdx* α7-integrin population accounted for the 6.8% of total events already in the non-perturbed condition; after cardiotoxin injury its number fluctuates around an average value of 7% of the total events with minor variations.

Once established the dynamic of participation of the MPs in the injury reaction, we shifted our attention on the fibroadipogenic compartment, constituted by Sca-1, CD34, CD140α, CD90.2 and Vimentin expressing cells. In wild type injured mice, the fibroadipogenic compartment, varies in time with a well-defined trend (Fig. 6c). In fact, similarly to the α7-integrin^+^ population, FAPs number increased reaching the maximum expansion at day 5. In contrast, mdx FAPs did not show any change as a response to cardiotoxin injury.

To conclude our analysis of the dynamic occurring after an injury, we also evaluated the abundance of vessel-associated populations whose function is strictly necessary to fully accomplish the regeneration process^28^.

These were identified among CD45 negative cells for their expression of Sca-1, CD31, CD146 and CD34 antigens, all markers of endothelial populations. Moreover, when considering α7-integrin antigen, we retraced its expression among the vessel-associated cells, indicating the inclusion of myoendothelial cell subpopulation.

In both the wild type and *mdx* models we could observe an initial down regulation of the endothelial population due to CTX injection in the first three days but after this, while the wild type cells increased to restore tissue integrity, the *mdx* vessel-associated cells did not (Fig. 6d).

## Discussion

The skeletal muscle tissue has a remarkable capability to self-repair after tissue damage. However, this healing process fails as a consequence of muscle related genetic disorders. For example, skeletal muscle dystrophies can impair the muscle repair process, leading to progressive muscle wasting and weakness characterized by chronic inflammation that eventually leads to the complete replacement of muscle fibers with fat and fibrotic deposition^29^. Ageing can also cause permanent changes in the regenerative potential of a healthy muscle. In fact, satellite cells (SCs) are affected by the ageing process due to changes in systemic or niche factors which lead to SC dysfunction and decline in number^12,13,30–33^.

Here we have exploited the resolution power of mass cytometry to characterize the changes of mononuclear cell population abundance under different muscle perturbation conditions, either chronic or acute. To this end we tested several antibodies and we ended up with a panel of 23 metal-tagged antibodies that can help to characterize the mixed mononuclear cell populations at single-cell level (Supplementary Table 1). In particular, we dwelt on muscle tissue ageing and on the cardiotoxin (CTX) injury-response in wild type and dystrophic muscles (*mdx*).

Our study showed that the number of mononuclear cell populations isolated from muscles of *mdx* mice was basically higher in comparison with wild type one (Supplementary Fig. 3a) during the lifespan of a year and a half considered in our experiments. The observed increase cannot be exclusively imputed to the inflammatory compartment, which is chronically activated in the dystrophic muscle tissue. In fact, the hematopoietic to non-hematopoietic cell ratio did not vary significantly in wild type and *mdx* mice (Fig. 3a). Nevertheless, the mechanical stress sensed from the muscle fibers induces a chronic state of inflammation as a result of the deteriorating environment^34,35^. The inflammatory cell number, as evaluated by the pan-macrophage F4/80 marker and including eosinophils, monocytes, M1 and M2 macrophages was higher when compared to the hardly detectable wild type inflammatory compartment (Fig. 3b). This is consistent with the notion that the dystrophic muscle is under chronic inflammation as a consequence of sarcolemma damage and fiber necrosis.

The availability of an antibody against the CD206 antigen allowed us to estimate the size of the pool of M2 macrophages. Consistent with their anti-inflammatory and pro-regeneration function, M2 macrophages were found to be less numerous at 1.5 and 8 months, in the regeneration window, while reaching their highest value at 3.5 months after birth during the degeneration phase (Fig. 3c)^36^. While our antibody panel did not include a reagent specific for M1 macrophages, it is important to note that the distinction between M1 and M2 macrophages is not clear-cut and evidence is accumulating that inflammatory infiltrates contain, in addition to a mixture of M1 and M2 macrophages, cells that share both M1 and M2 properties^17,19,37^.

We also evaluated α7-integrin^+^ cell abundance, to estimate the number of cells in the myogenic compartment. Since we did not have a reliable reagent for Pax7, the satellite cell marker of choice, we estimated the number of satellite cells by looking at CD45^−^, Sca-1^−^, α-7integrin^+^, CXCR4^+^ and CD34^low^. We could not observe significant differences in this population with myogenic potential both in wild type and *mdx* mice in all the conditions tested (Fig. 3d). Perhaps unexpectedly, we did not observe a decrement in time of α7-integrin^+^ cells in both the wild type and *mdx* models and we were not able to observe the satellite cell depletion associated to the ageing or the dystrophic muscle tissue.

Nevertheless, the high dimensional analysis, made possible by mass cytometry, revealed that wild type and *mdx* α7-integrin^+^ cells differ in the expression of the CXCR4 antigen. CXCR4 is the receptor for the chemokine stromal-derived factor 1 (SDF-1) which has been reported to be secreted in the muscle tissue and to be implicated in different cell processes such as chemotaxis, proliferation, differentiation and calcium flux^24^. In particular, we observed that the α7-integrin^+^ cells derived from *mdx* muscles contained a higher fraction of cells expressing the CXCR4^+^ antigen (Fig. 4a). This observation lead us to speculate on a possible role of the SDF-1/CXCR4 crosstalk in the dystrophic pathology^12–14,38,39^.

Fibro-adipogenic progenitors (FAPs) have been suggested, to play a pivotal role^15^ in the progressive substitution of muscle tissue with fat and fibrotic scars, which contributes to exacerbate the dystrophic phenotype. In our study, we monitored FAPs as Sca-1^+^/CD140α^+^/CD90.2^+^/CD34^+^ cells. As anticipated, the number of FAPs was on average 3-fold higher in *mdx* mice and after reaching a peak in 8 month old *mdx* mice, it rapidly decreased, possibly as a consequence of the attenuation of the inflammation process (Fig. 3f). The antibody panel used in these experiments also included some reagents against activated intracellular regulatory proteins. We observed that *mdx* FAP cells were highly positive for the expression of the phosphorylated form of the transcription factor Creb (Fig. 4e). Creb is activated when phosphorylated and in this form acts as a master regulator of adipogenesis^25^. This observation is consistent with the involvement of FAPs in fat deposition in the muscles of dystrophic mice and suggests that their adipogenic commitment already starts at 1.5 months after birth.

We also monitored the expression of interleukin-6 (IL-6) in fibroadipogenic progenitors based on the evidence that FAPs are a damage-inducible source of the IL-6 chemokine^18^.

In *mdx* FAPs we observed a well-defined trend in FAPs IL-6 production, which contributes to exacerbate the dystrophic muscle phenotype, by sustaining the inflammatory response and the repeated cycles of muscle degeneration and regeneration (Fig. 4d).

Our hypothesis is that the observed bursts in IL-6 production in *mdx* FAPs from 1.5 and 8 month old mice are linked to the regeneration phase of the myofibers, since literature reports that IL-6 chemokine production contributes to muscle regeneration after injury and this may have a stimulatory effect on satellite cells^18^.

Our analysis on the FAP cell population also highlighted a precise balance in Sca-1 expression, which is furthermore closely correlated to CD34 expression (Fig. 4b). In particular, we observed that the number of cells expressing Sca-1 (or CD34) was higher in *mdx* mice in all the conditions but that the ratio between Sca-1-CD34^low^/Sca-1-CD34^high^ expressing cells was maintained over the mouse lifespan (Fig. 4c).

Since it has been reported that the Sca-1 marker is required for the self-renewal of mesenchymal stem cells^40^, while CD34 is related to cell proliferation^41^, we asked ourselves if this observed trend could have an important role in ensuring correct tissue homeostasis.

In addition, we must mention the identification of a population with a well-defined trend during ageing (Fig. 3g) and a clearly outlined expression profile (Fig. 3e), even if it does not correspond to any known population in literature. This new population was characterized by a high expression of Sca-1 and CD34, as also observed in the “classical” fibroadipogenic progenitors, but it is distinct from FAPs for the absence of expression of two additional FAPs markers, CD140α and CD90.2 (Fig. 3e).

We also identified a second rare population, never described before representing approximately 1.6% of the total mononuclear population of wild type and *mdx* ageing muscles. This population was positive for the expression of CD146, CD90.2 and CD140β, markers that are typical of pericytes^42,43^, but at the same time it was also positive for α7-integrin whose expression was never associated to pericytes before (Fig. 2d).

We also examined a model of acute damage/regeneration obtained by injecting into the muscle the snake venom cardiotoxin, which represents a well-established model of muscle damage and regeneration.

By inducing the irreversible depolarization and contraction of the myofibers, cardiotoxin triggers necrosis, inflammation and sarcolemma disruption, thus recapitulating all the phases commonly accompanying muscle injury^44,45^.

The regeneration process was monitored by analyzing samples at the mass cytometer at days 1, 3, 5, 10 and 20 after cardiotoxin injection of wild type or *mdx* mice.

A time-dependent variation in the abundance of muscle total mononuclear cells (Supplementary Fig. 3b) was observed for wild type mice, indicating that the system suffered the damage but promptly responded (already at day 3) to restore muscle tissue homeostasis. The maximum peak in the total cell number was observed 3 days after CTX injury and corresponds to the peak of the CD45^+^F4/80^+^ hematopoietic cells (Supplementary Fig. 3b, 3 days after the injury). Inflammatory cells were the predominant population three days after the injury, representing up to 30% of the total mononuclear cells (Fig. 6a). The activation of the inflammatory compartment is limited to the first few days after the injury as we observed that the number of macrophages returns to almost baseline at day 10.

Fibroadipogenic progenitors, here identified as Sca-1^+^, CD34^+^, CD140α^+^ and CD90.2^+^ stimulate satellite cell activation and differentiation, thus playing a positive role in muscle regeneration^18^. FAPs are quiescent in intact muscles but efficiently proliferate in response to injury^16,18,19^. Consistently we observe that FAPs rapidly expanded between day 3 and 5, returning to baseline level once finalized their function of providing a transient production of pro-differentiation signals for the myogenic progenitors and of depositing extracellular matrix (Fig. 6c).

In response to the secreted inflammatory and fibroadipogenic stimulating signals, the myogenic compartment promptly activates itself. After an initial decrease, α7-integrin^+^ cells, which include satellite cells and myoblasts (Fig. 6b), start to expand at day 3, reaching their maximum at day 10 when, following the attenuation of the inflammatory response and FAP expansion, decrease in number. The coordination of these processes confirms the importance of this three-way cross-talk for a correct regeneration process.

A different kinetics is observed for endothelial progenitor cells, here characterized as CD31, CD34 and Sca-1 expressing cells. The reconstruction of the blood vessel network only takes place at the end of the regeneration process once inflammatory cells have been removed and FAPs have decreased in number (Fig. 6c, d).

Wild type regeneration after acute injury occurs in a well-defined and coordinated manner to ensure the correct restoration of skeletal muscle tissue and to avoid the unpleasant consequence deriving from its irreparable impairment.

We also investigated the response to acute injury of a muscle environment that is already chronically perturbed as in *mdx* mice. The total mononuclear cells extracted from muscle of *mdx* mice (Supplementary Fig. 3b) did not show any significant increase following cardiotoxin injury, thus indicating that the system does not respond to cardiotoxin injury through population expansion or, possibly, that the increase of a population is counterbalanced by a decrease in another one, in a way that the total cell number is not altered. We favor the latter hypothesis since we observed that after three days the *mdx* inflammatory cells significantly expand (Fig. 6a) while the endothelial cell number decreases (Fig. 6d). The myogenic and fibroadipogenic compartments, on the other hand remain rather constant without significant variations during the regeneration process (Fig. 6b, c).

Overall, our multiparametric analysis offers a comprehensive description of both muscle tissue homeostasis and the rearrangements in population abundance induced by a perturbation of the muscle system, be it a chronic one in *mdx* mice or an acute one after cardiotoxin injury.

## Methods

### Animal procedures

Wild type C57BL/6 or C57BL/10ScSn-*Dmd*^mdx^/J (also known as mdx) mice were purchased from the Jackson Laboratories. Mice were bred respecting the standard animal facility procedures and experiments on animals were conducted according to the Animal Research Ethical Committee in compliance with the Italian Ministry of Health.

Wild type or *mdx* mice were employed at the specified age (1.5, 3.5, 8 and 16 months after birth for ageing muscles experiments and 1.5 for muscle injury ones) and cervical dislocation (CD) euthanasia was performed using appropriate equipment and in accordance with the guidelines. Experiments on animals were conducted according to the rules of good animal experimentation I.A.C.U.C. n°432 of March 12 2006.

### Muscle injury experiments

Hind limb muscles (tibialis anterior, quadriceps and gastrocnemius) of 1.5 month old C57BL/6 and C57BL/6ScSn-Dmdmdx/J mice were injected with 10 μM cardiotoxin (Latoxan L81-02) under anesthesia performed with an intramuscular injection of physiologic saline (10 ml/Kg) containing ketamine (5 mg/ml) and xylazine (1 mg/ml). The animal care and experimental plan were conducted according to the rules of good animal experimentation I.A.C.U.C. n°432 of March 12 2006 and under ethical approval released on 16/09/2011 from The Italian Ministry of Health, protocol #163/2011-B.).

Treated mice were all sacrificed at 20 days after injury, considering the time points of 1, 3, 5, 10 and 20 days (three mice per time point). Non-injured muscles were used as controls.

### Skeletal muscle mononuclear cell isolation

For the dissociation of muscle tissue and isolation of single mononuclear cells, mice hind limbs were dissected and finely minced mechanically with tweezers. Then the small muscle pieces obtained were subjected to an enzymatic digestion for 1h at 37°C performed using 2 μg/ml collagenase A (Roche cat#10103586001), 2,4U/ml dispase II (Roche cat#04942078001) and 0,01mg/ml DNase I (Roche cat#04716728001) diluted in D-PBS with Calcium and Magnesium (Gibco, cat#14040).

Once digested, the enzymatic reaction was stopped with Hank’s Balanced Salt Solution with calcium and magnesium (HBSS Gibco, cat#14025-092) supplemented with 0.2% BSA and 1% Penicillin-Streptomycin (P/S, 10.000 U/ml). The resulting cell suspension was subjected to sequential filtration through 100 μm, 70 μm and 40 μm cell strainers and centrifugations at 600 x g for 5 min.

### Skeletal muscle fixation and immunohistological analysis (H&E and Sirius red staining)

Isolated tibialis anterior (TA) muscles were collected and snap frozen in liquid-nitrogen, embedded in OCT for transverse cryo-sectioning with a Leica cryostat. Cryosections (10 μm thickness) were stained with hematoxylin and eosin (H&E) or Sirius red (Sigma-Aldrich) according to the analysis carried out.

As regarding H&E, cryosections were fixed with 4% paraformaldehyde (PFA) for 15 minutes at room temperature (RT). After washing in pure water, sections were incubated in the hematoxylin solution for 15 minutes at RT and rinsed for 5 minutes in tap water. After washing in pure water, sections were counterstained with an alcoholic solution of eosin for 30 minutes at RT. Following the eosin staining, sections were rinsed, dehydrated in growing concentrations of alcohol, treated with a clearing solution (Histo-Clear, Agar Scientific) and finally mounted, using the resinous Eukitt mounting medium (EMS Catalog #15320).

For the Sirius Red staining, cryosections were fixed in Bouin solution for 1 h at 56° C, then washed with running tap water and incubated for 1 h at room temperature with a solution containing 0.1% Direct Red 80 (Sigma-Aldrich, cat#365548) in a saturated aqueous solution of picric acid (Sigma-Aldrich, cat# P6744-1GA). Then, they were washed in two changes of acidified water, dehydrated in three changes of 100% ethanol and cleared in xylene. Cryosections were finally mounted, using the resinous Eukitt mounting medium. H&E and Sirius Red stained sections were photographed on a Zeiss Lab A1 AX10 microscope.

### RNA isolation, reverse transcription (RT) and real-time quantitative PCR

Total RNA was isolated from muscle tissue using NucleoZOL reagent (Macherey-Nagel, cat#740404.200) according to manufacturer’s recommendations. cDNA was synthesized from 1 μg of total RNA with PrimeScript RT Reagent Kit (Takara, cat#RR037A). RT-PCR was performed by using SYBR Premix Ex Taq (Tli RNaseH Plus) (Takara, cat#RR820Q) with 10 μM each primer and normalized to *Actin* as a housekeeping gene.

Primer list:

***Actb***

Fw: 5′-CACACCCGCCACCAGTTCGC-3′

Rv: 5′-TTGCACATGCCGGAGCCGTT-3′

***Colla1***

Fw: 5′-GCATTCACCTTTGAAACTTAGT-3′

Rv: 5′-CTTCAAGCAAGAGGACCAA-3′

***Col3a1***

Fw: 5′-CAACGGTCATACTCATTC-3′

Rv: 5′-TATAGTCTTCAGGTCTCAG-3′

## Mass cytometry

### Staining

3x10^6^ cells from each sample were used for the mass cytometry staining. Cells were centrifuged at 600 x g for 5 min and washed in D-PBS w/o Calcium and Magensium (BioWest cat#L0615-500). Dead cells were stained by resuspending the samples in D-PBS with cisplatin at a final concentration of 5 μM and by incubating them for 5 min at room temperature. The reaction was quenched with 2 ml of MaxPar Cell staining buffer (Fluidigm, cat#201068) and the samples were centrifuged at 600 x g for 5 min. The cells were fixed with 1 mL Fix I Buffer (Fluidigm, cat# 201065) and incubated for 10 minutes at room temperature. The samples were then centrifuged at 800 x g for 5 min and washed twice with 1 mL of Barcode Perm Buffer (Fluidigm, cat# 201057). Each sample was then resuspendend in 800 μL Barcode Perm Buffer while barcode dyes were resuspendend completely in 100 μL of Barcode Perm Buffer and transferred to the appropriate sample. The samples were mixed completely and incubated for 30 min at room temperature. Cells were centrifuged and washed twice with MaxPar Cell Staining Buffer. Each sample was then resuspendend in 100 μL MaxPar Cell Staining Buffer and all the barcoded ones were collected into one unique tube that was subjected to the antibody staining protocol. Cells were centrifuged and incubated with antibodies against surface markers included in the used mass cytometry panel for 30 min at room temperature. Cells were washed twice by adding MaxPar Cell Staining Buffer, centrifuged 800 x g for 5 min and permeabilized with methanol for 10 min on ice. Cells were washed twice with MaxPar Cell Staining Buffer and stained with antibodies against intracellular markers included in the mass cytometry panel for 30 min at room temperature. Cells were washed twice with MaxPar Cell Staining Buffer and stained for 1 h at room temperature with the intercalation solution, composed of Cell-ID Intercalator-Ir (Fluidigm, cat#201192A, 125 μM) into MaxPar Fix and Perm Buffer (Fluidigm, cat#201067) at a final concentration of 125 nM (1000x dilution of the 125 μM stock solution). Cells were washed twice with MaxPar Cell Staining Buffer and MaxPar Water supplemented with 0,1% Tween respectively and kept in a pellet on ice until measured.

### Acquisition of single cell

Just before the run of the samples, cells were re-suspended in 10% EQ Four Element Calibration Beads (Fluidigm, cat# 201078) and acquired on the CyTOF2™ mass cytometer (Fluidigm; DVS Sciences). The instrument was tuned and mass calibrated according to manufacturer’s instructions. During the acquisition the data was calibrated by dual-count calibration on the instrument. Data was then converted on the fly to fcs files using the manufacturer’s settings.

### Antibody panel

In studying the literature and previous works, a list of antigens was compiled and used for single-cell CyTOF2 analysis. The full list of the used antibodies to discriminate the main cell populations and their activation state, are listed in Supplementary Table 1.

### Data processing

Following data acquisition, channel intensity was normalized using calibration beads^46^. Data was gated using CytoBank^47^.

### Statistical analysis

Data were from at least 3 independent samples, unless otherwise indicated. Results are presented as means ± SEM. Statistical evaluation was conducted by using the two-way ANOVA. Comparisons were statistically considered significant at statistical significance defined as * p < 0.05; ** p < 0.01; *** p < 0.001; **** p < 0.0001. All statistical analysis was performed using Prism 7 (GraphPad).

